# High-performance Flexible Microelectrode Array with PEDOT:PSS Coated 3D Micro-cones for Electromyographic Recording

**DOI:** 10.1101/2022.02.03.479004

**Authors:** Jiaao Lu, Muneeb Zia, Matthew J. Williams, Amanda L. Jacob, Bryce Chung, Samuel J. Sober, Muhannad S. Bakir

## Abstract

High signal-to-noise ratio (SNR) electromyography (EMG) recordings are essential for identifying and analyzing single motor unit activity. While high-density electrodes allow for greater spatial resolution, the smaller electrode area translates to a higher impedance and lower SNR. In this study, we developed an implantable and flexible 3D microelectrode array (MEA) with low impedance that enables high-quality EMG recording. With polyimide micro-cones realized by standard photolithography process and PEDOT:PSS coating, this design can increase effective surface area by up to 250% and significantly improve electrical performance for electrode sites with various geometric surface areas, where the electrode impedance is at most improved by 99.3%. Acute EMG activity from mice was recorded by implanting the electrodes *in vivo*, and we were able to detect multiple individual motor units simultaneously and with high resolution (SNR ≫ 100). The charge storage capacity was measured to be 34.2 mC/cm^2^, indicating suitability of the electrodes for stimulation applications as well.

## I. Introduction

Compared with central nervous system recording, electromyographic recordings (EMGs) provide more precise and subtle information about muscle movements, that could be critical for specific motor functions characterization and behavioral disease progression [1]. Traditionally, EMGs are collected by inserting fine wires directly into the muscles [2]. However, fine-wire electrodes predominantly capture bulk EMG consisting of activity of multiple units, whereas single motor units are essential for holistic computational analyses [3], [4]. To acquire single motor unit EMG, two key enablers are low electrode impedance and high spatial resolution. Both are important for analysis methods in neuroscience because low impedance ensures that the signal-to-noise ratio is high so that neural events can be detected and high spatial resolution provides information across different recording sites that is needed to reliably identify events that are generated by the same source. However, these metrics are inherently antagonistic: high spatial resolution demands smaller, densely packed electrode sites whereas electrode impedance is negatively impacted by reduced electrode surface area [5].

One way to address this challenge is to increase the effective surface area (ESA) while maintaining the geometric surface area (footprint). Previous techniques attempting to achieve high ESA for better electrochemical performance can be roughly categorized in two ways: surface modification and 3D structure fabrication. As for surface modification, coating electrode sites with porous conductive polymer [6], [7], or plasma treatment of the metal surface [8] are some of the ways that have been used to increase the ESA and electrical performance of the electrodes. Fabricating 3D electrodes is another way to increase ESA, however, previously reported methods are either difficult to scale to higher densities or they require relatively complex fabrication processes that are challenging to be adapted on flexible substrate [9]–[13]. Combining surface modification and scalable 3D structure fabrication, this paper aims to achieve high-density microelectrode array (MEA) that is flexible and bio-compatible for *in vivo* application.

In this work, the 3D electrode sites are fabricated using polyimide micro-cones that are subsequently coated with poly(3,4-ethylenedioxythiophene) polystyrene sulfonate (PEDOT:PSS) to achieve low impedance. We demonstrate the improvement of electrical performance by comparing the impedance between 2D and 3D electrodes with different finishing materials, and show that with the proposed design, the impedance is improved by 99.3% when compared to the planar gold electrode. *In vivo* data of the recorded EMG from mouse digastric muscle is also presented; from the measured EMG a high signal-to-noise ratio (SNR) of 447.3 was calculated.

## II. Fabrication of 3D MEAs

The fabrication process of 3D microelectrode arrays (MEAs) is summarized in Fig. 1. At first, a 15-μm thick polyimide (PI-2611) layer was spun and cured on a 4-in silicon carrier wafer (Fig. 1(a)). To build 3D polyimide micro-cones, a 15-μm layer of photodefinable polyimide (HD-8820) was spun, soft baked and patterned by a standard photolithography process in a maskless aligner (Fig. 1(b)). The patterned 3D micro-cones were subsequently cured in oven. A reactive ion etch (RIE) process using O_2_ and SF_6_ (5:1 ratio) plasma was performed to roughen the surface of polyimide substrate and cones. This further increases the surface area and improves the metal adhesion on polyimide. The traces and electrodes were patterned using a lift-off process; a titanium (Ti) adhesion layer followed by gold (Au) were deposited using evaporation process. A thin sacrificial Ti layer was also deposited on the gold as a means to visually identify successful etch of the top polyimide layer. (Fig. 1(c)). A 5-μm polyimide (PI-2611) was then spin coated and cured in oven. An etch mask was patterned using a 6-μm thick positive photoresist (PR) and the top polyimide was then etched using RIE to expose the electrode and connector sites (Fig. 1(d)). The MEAs were subsequently diced using a femtosecond laser (Fig. 1(e)). A 32-channel Omnetics connector was then connected to each array using anisotropic conductive film (ACF) (Fig. 1(f)).

**Fig. 1.**
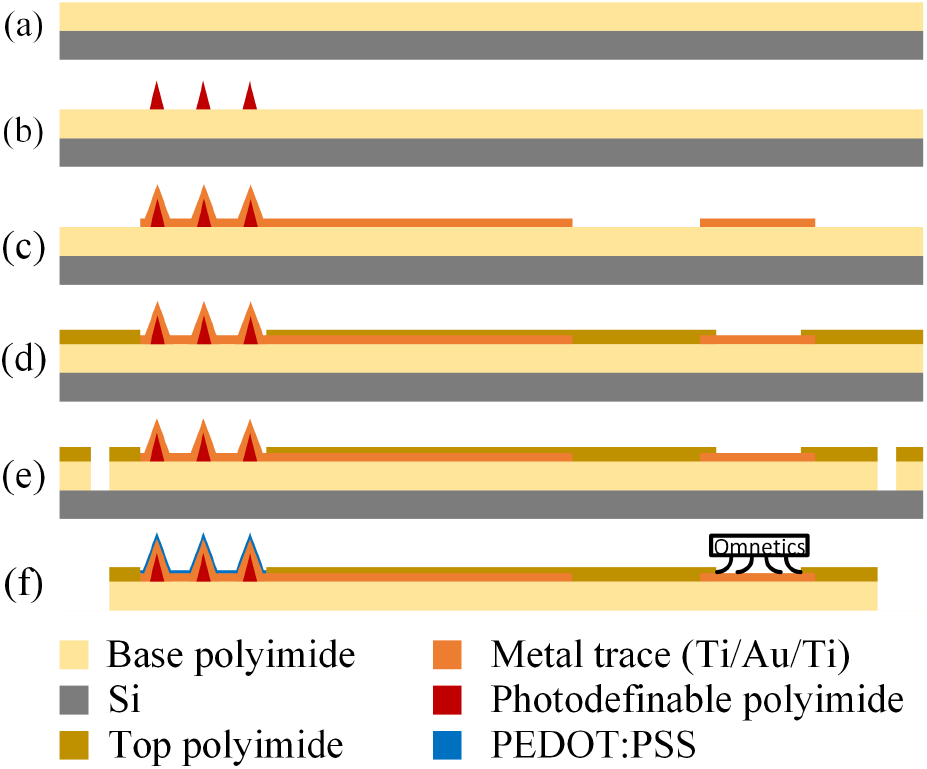
Fabrication process of the flexible 3D microelectrode arrays. (a) Base polyimide coating. (b) Photodefinable polyimide coating and patterning. (c) Ti/Au/Ti evaporation and patterning. (d) Top polyimide coating and etching. (e) Laser dicing. (f) Device release, assembly and electropolymerization of PEDOT:PSS.

To further improve the electrical performance, a PEDOT:PSS layer was deposited onto the gold microelectrodes via electropolymerization [6]. An aqueous dispersion of 10mM 3,4-Ethylenedioxythiophene (EDOT) added to 2.0g/100mL sodium polystyrene sulfonate (NaPSS) was used as electrolyte to electropolymerize PEDOT:PSS under galvanostatic conditions in a two-electrode setup, where the gold electrode array served as working electrode and a platinum mesh as counter/reference electrode. A constant current of current density 0.5 mA/cm^2^ was applied for 5 min to deposit a layer of PEDOT:PSS under room temperature.

The PEDOT:PSS coated 3D polyimide micro-cones were successfully fabricated in electrodes with various footprint sizes (from 200μm×200μm to 5μm×5μm). An optical image of fabricated MEA is shown in Fig. 2(a). The surface morphology of PEDOT:PSS covered cones was examined with a field emission scanning electron microscope (SEM), and an example of 20μm× 20μm electrode is shown in Fig. 2(b). The conical shape of the 3D micro-structure is achieved by tuning the exposure and development process for the photodefinable polyimide. The addition of 3D micro-cones results in an increase in ESA by 150% to 250% for different footprint sizes. Electrodeposited PEDOT:PSS uniformly covered the 3D micro-cones, forming a rough and porous coating to further enhance the ESA. The crosssectional view in Fig. 2(c) clearly reveals the different layers of the electrode: the substrate and cone made of polyimide, thin trace metal, and the top PEDOT of the thickness in the order of 1 μm.

**Fig. 2.**
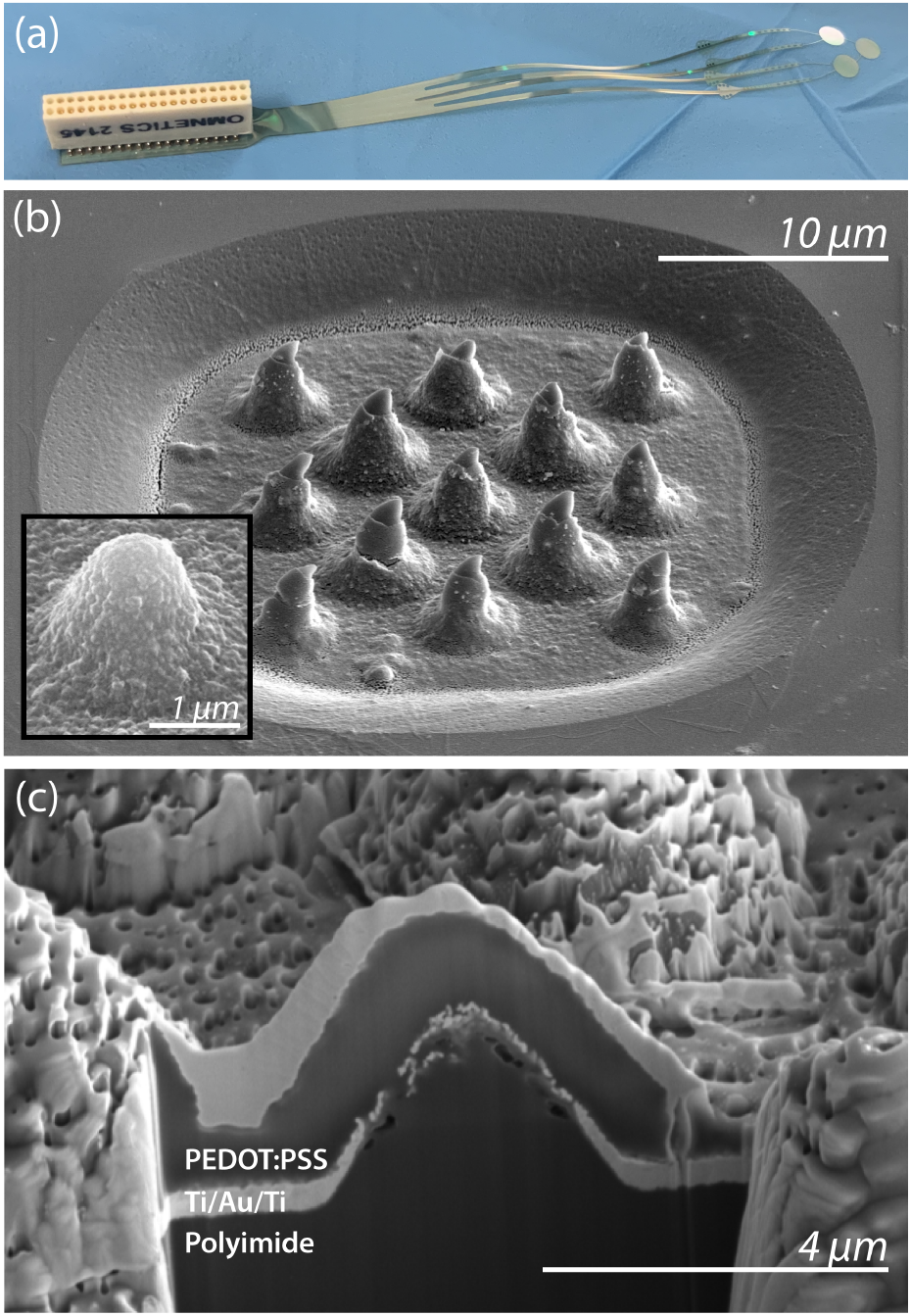
Optical and electron microscopy. (a) An optical image of the fabricated 3D MEA. (b) A SEM image of a 20μm×20μm electrode pad with 13 cones on top to increase the effective surface area. The inset picture shows the surface contains nano features and porosity after PEDOT:PSS electropolymerization. (c) The SEM of focused ion beam (FIB) cross-section of the electrode site showing multiple layers of the device. A thin layer of tungsten was deposited on top to protect the surface during FIB.

## III. Characterization and EMG measurement

### A. Electrochemical characterization

In order to evaluate the improvement in electrical performance from the 3D structure and porous PEDOT:PSS coating, impedance at 1kHz of planar and 3D electrodes both with gold and PEDOT:PSS finishing layers were obtained in a 2-electrode setup in physiological saline solution (0.9% NaCl) using a Ag/AgCl reference electrode. We compared the average impedance (n=8) of electrode sites across eight different sizes as shown in Fig. 3. Electrodes with 3D micro-cones show a significant impedance reduction across all footprint sizes for both gold and PEDOT:PSS surface finishes. When compared to the gold electrodes, the PEDOT:PSS treatment further reduces the impedance of the 3D micro-cone electrodes which aligns with previous observation [6]. For 10μm×10μm recording sites, there was a 99.3 % reduction in impedance of the PEDOT:PSS coated 3D micro-cone electrodes as compared to the planar gold electrodes. We are able to achieve high-density recording sites using the 3D micro-cone structures as the average impedance of the smallest test size for 3D micro-cones (5μm×5μm) was 146.5 kOhm, which was the same as the impedance for a planar gold electrode that was ~250 times larger (80μm×80μm).

**Fig. 3.**
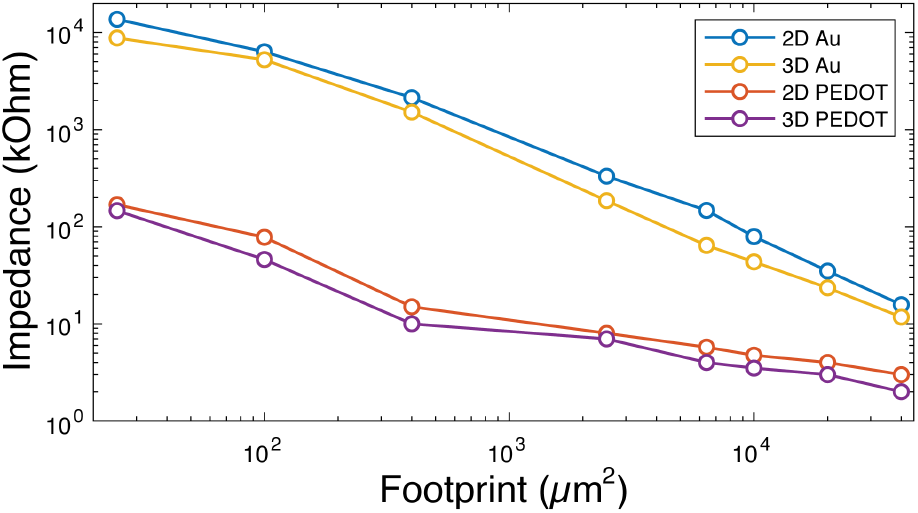
Average impedance (n=8) at 1kHz across eight footprint sizes for planar and 3D electrodes with gold and PEDOT:PSS finishing layers.

We further validated the decrease of impedance by comparing the electrochemical impedance spectroscopy (EIS) results between planar gold and 3D PEDOT:PSS coated micro-cone electrodes with representative footprint sizes, as shown in Fig. 4. The experiments were conducted in saline solution with a 3-electrode system with Ag/AgCl reference and platinum mesh counter electrodes. The changes in impedance magnitude are consistent with our results at 1 kHz, as the 3D micro-cones together with PEDOT coating significantly reduce the impedance value over the tested frequency range from 0.1 Hz to 100 kHz for all sizes.

**Fig. 4.**
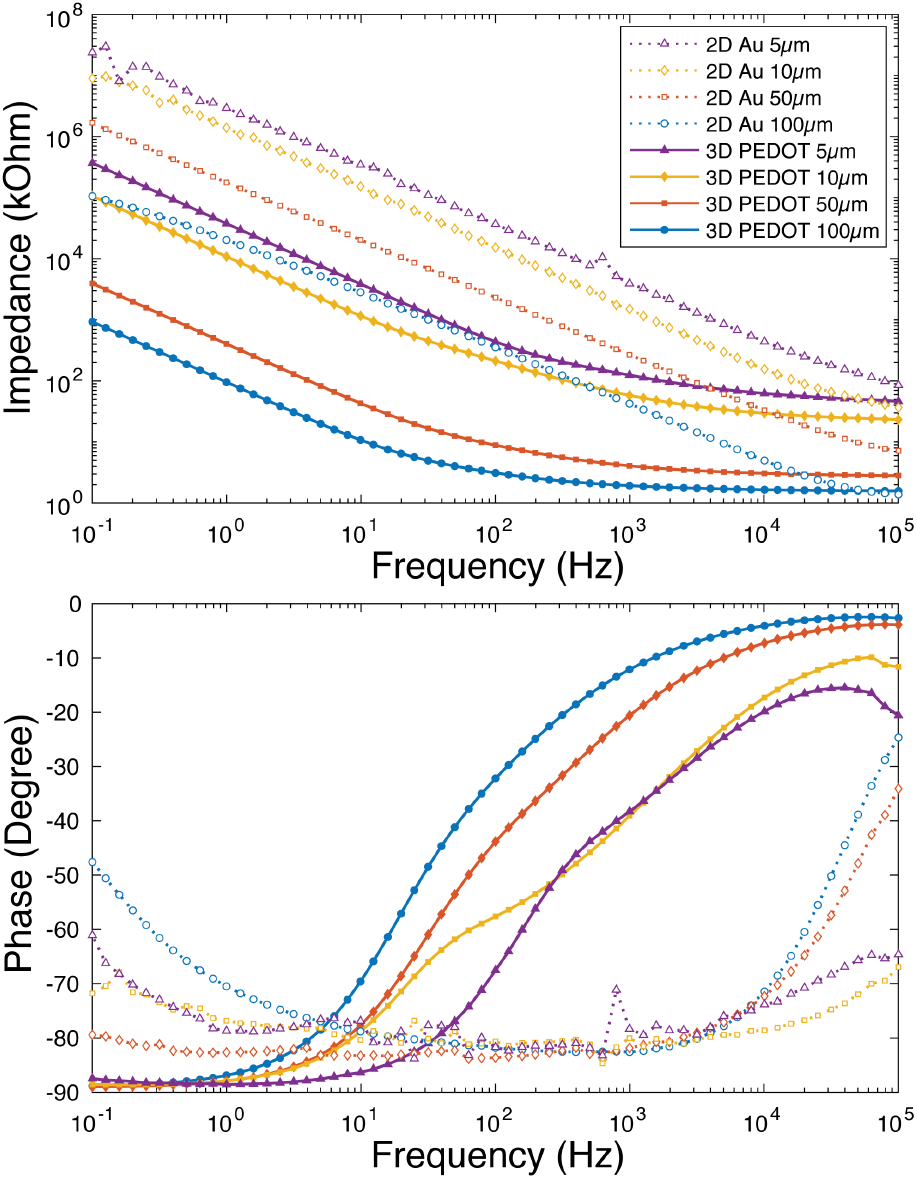
EIS measurement of 3D PEDOT:PSS coated microelectrodes and 2D gold microelectrodes with four different footprint sizes. Legend applies to both graphs; the number values correspond to the side lengths of the square electrode site.

Cyclic voltammetry (CV) measurements were also carried out with the same 3-electrode setup to determine the charge storage capacity (CSC) of the fabricated 3D micro-cone electrodes. The CV results of 3D micro-cone electrode with footprint of 200μm×200μm is shown in Fig. 5 and the CSC is calculated to be 34.2 mC/cm^2^. This demonstrates its feasibility for stimulation application with the CSC comparable or better than some of the published work [14], [15].

**Fig. 5.**
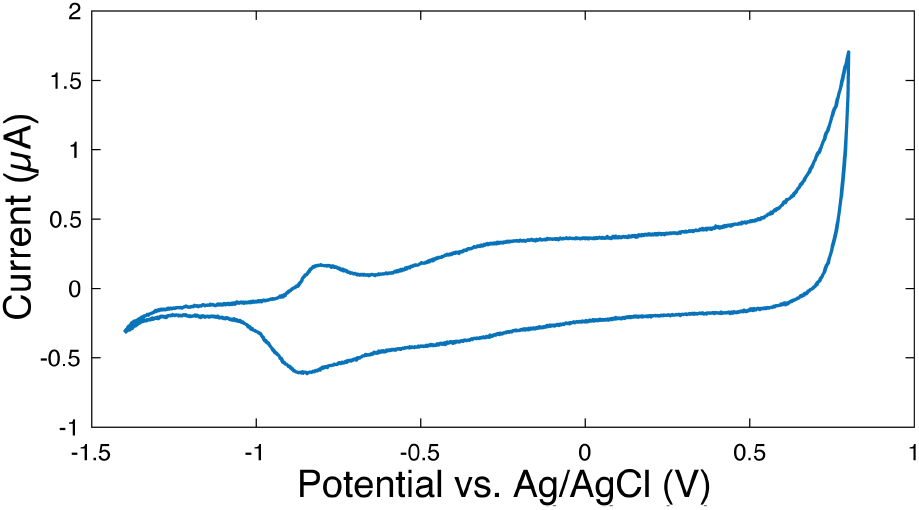
Cyclic Voltammetry of PEDOT:PSS coated 3D micro-cone electrode with footprint size of 200μm×200μm in saline solution, swept at 100 mV/s.

### B. EMG measurement and data analysis

The fabricated 3D MEAs with micro-cones were utilized to record EMG activity from digastric muscles of anesthetized mice. All procedures were approved by the Emory University Institutional Animal Care and Use Committee. Anesthesia was induced in 2 adult C57BL/6J male mice using 4% vol/vol isoflurane in oxygen gas and maintained at appropriate levels throughout the procedure and data collection. An incision was made under the jaw, and skin was removed to reveal the digastric muscle. The electrode array was placed on the muscle’s surface to collect spontaneously-occurring motor unit activity that was amplified via a 32-channel digital amplifier headstage (RHD2132; Intan Technologies). EMG signals were recorded on a computer at 30 kHz through a RHD USB Interface Board (Part C3100; Intan Technologies).

Waveforms from different motor units were identified using standard methods that use principal component analysis (PCA) and k-means to cluster waveforms with similar shapes [16]. The amplitude of each spike cluster was measured by the average peak of each waveform in that cluster. The noise level of each channel was measured by the root mean square (RMS) of a one-minute section of signal not containing any spikes. The SNR for each spike was then computed by dividing the amplitude of each spike cluster by the noise level on the corresponding channel.

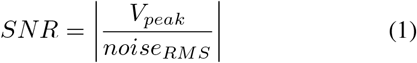

High-quality EMG recordings were successfully obtained from mice with the low-impedance 3D MEAs, in which both single and multiple motor units were observed (Fig. 6). In single motor unit recording, the device were able to detect small differences in the action potential’s timing and amplitude, which indicate that the electrode has high temporal and spatial resolution to distinguish the different locations of electrode contacts relative to the active muscle fibers. These variations across channels allow multiple motor units to be isolated from single recordings. The multiple motor units recording further showed that our device has significantly higher SNR compared to previously published work [11]. We achieved a mean SNR value of **203.3** for the motor unit with the smaller spike, as well as a mean SNR of **447.3** for the motor unit with the larger spike (Fig. 6(b) top).

**Fig. 6.**
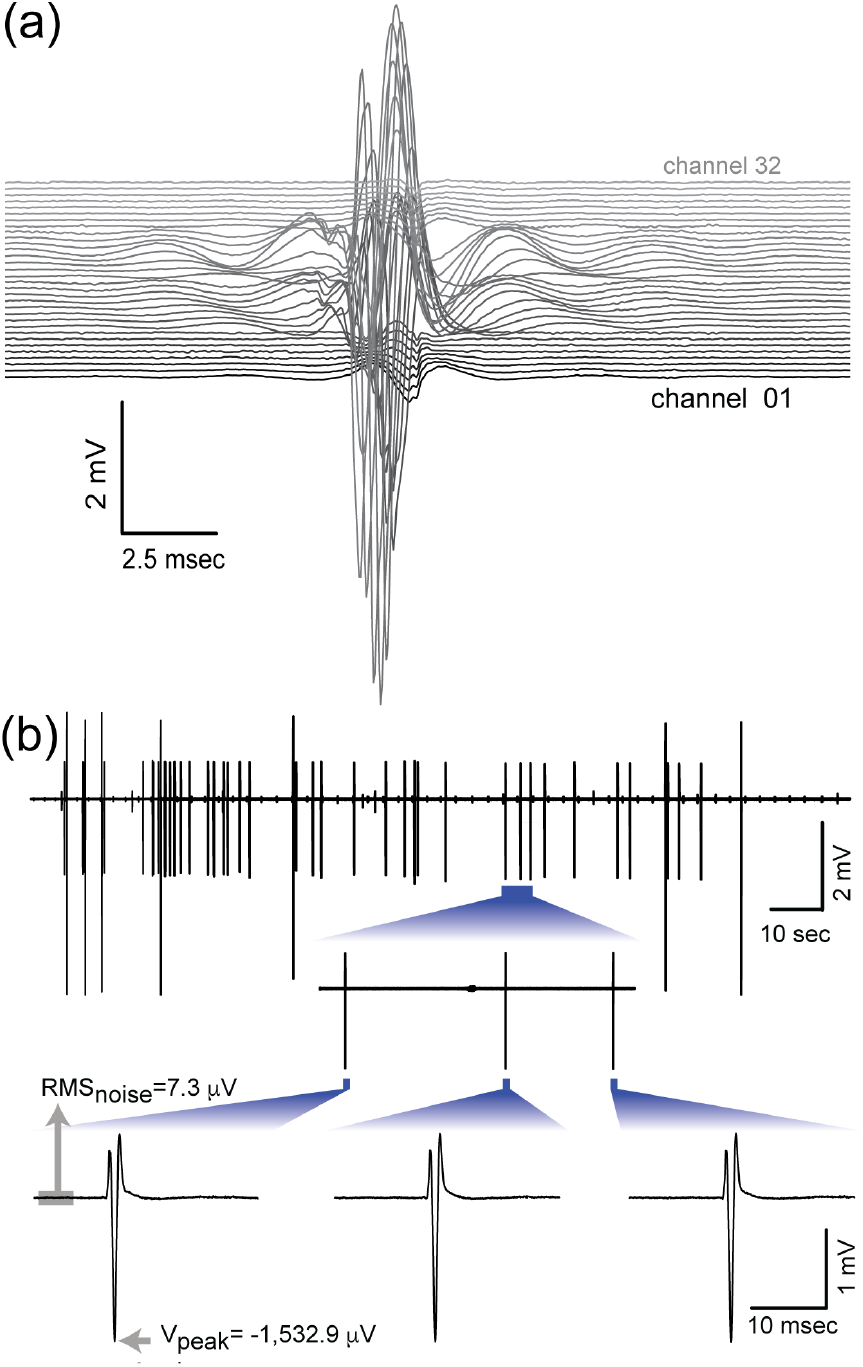
High-resolution EMG recordings with micro-cone MEAs. (a) Example recording of a single motor unit from a mouse digastric muscle. Here, a single action potential is detected by roughly half of the electrode array’s 32 channels. (b) Recording of multiple motor units from a single MEA channel. Top, recording from the digastric muscle during an epoch in which two different motor units distinguished by different amplitudes were recorded simultaneously. Bottom, detail of three individual smaller spikes and their corresponding peak voltage (V_peak_) and noise level (RMS_noise_).

## IV. Conclusion

In this paper, we present a flexible and bio-compatible microelectrode array with 3D micro-cones and PEDOT:PSS coating that gives a high SNR in *in vivo* experiments. Through the proposed novel fabrication process, we built micro-cones with photodefinable polyimide that increases the effective surface area up to 250%. Electrochemical characterizations were performed for electrodes with various sizes, where the impedance of our device consistently outperform benchmark devices in tested frequency range. Specifically, the impedance at 1kHz is reduced by two orders of magnitude. The cyclic voltammetry results show a CSC value of 34.2 mC/cm^2^, which suggest our device as a good candidate for stimulation applications. The 3D micro-cone electrodes were successfully implanted in mouse digastric muscles to obtain acute data, where a SNR up to 447.3 was measured through multiple motor units recording. With the high SNR achieved, our device provides a promising approach for the successful acquisition and analysis of single motor unit activity.

## Acknowledgment

This work was funded by National Institutes of Health Grant R01 NS109237, the McKnight Foundation and the Simons Foundation as part of the Simons-Emory International Consortium on Motor Control, and was performed in part at the Georgia Tech Institute for Electronics and Nanotechnology.

